# Three-dimensional morphological analysis of the dynamic digestive system in the green brittle star

**DOI:** 10.1101/640235

**Authors:** Daiki Wakita, Keisuke Naniwa, Hitoshi Aonuma

## Abstract

Brittle stars (Echinodermata: Ophiuroidea) digest a great diversity of food in their stomach, which widely lies in the central disk. As for a possible digestive activity, the green brittle star *Ophiarachna incrassata* is known to show a dynamic movement at the disk. This phenomenon would deeply involve the morphological structure of the stomach. However, past anatomical studies have shown the digestive system in two dimensions after wide incision of the body wall anchoring the stomach. This methodology restrains us from understanding how the stomach actually shapes inside a brittle star. We aim to visualize the morphology of brittle stars’ digestive system in a non-destructive and three-dimensional way, with a comparison between a relaxed specimen and a specimen fixed at the very moment of the disk’s movement. Employing X-ray micro-computed tomography (micro-CT) and introducing an instant freezing method with cryogenic ethanol, we found the stomach wholly transformed during the movement. We here brought transparency to the *in vivo* position of gut contents to hint the mechanism and digestive function of the movement. Our outcome spotlights a dynamic digestive process in echinoderms and a widely applicable method for probing into its relation with body structure.

## Introduction

More than 2,300 species of brittle stars (Echinodermata: Ophiuroidea) are known worldwide and constitute the largest class among extant echinoderms (Stöhr et al. 2019). Diversity in feeding habits could be an explanatory factor for the current success of this group (Fontaine, 1965). Their food varies in sort and scale, from sediment or small organisms including diatoms, dinoflagellates, foraminifera, and copepods, to the whole or part of large organisms such as polychaetes, bivalves, crabs, fish, other echinoderms, and sessile algae (Nagabhushanam & Colman, 1959; Fontaine, 1965; Hendler & Miller, 1984; Pearson & Gage, 1984; Ambrose, 1993). Their radially extending arms play a role in capturing food (Fontaine, 1965; Warner, 1971; Reimer & Reimer, 1975; Hendler & Miller, 1984), and then internal digestive organs take center stage.

Contrary to the various targets in feeding, brittle stars share the general structure of the digestive system. Its mouth is followed in order by buccal cavity, pharynx, esophagus, and stomach (Schechter & Lucero, 1968). Lacking intestine and anus, it terminates with the stomach, which occupies a large space inside the disk—central part of the body (Smith, 1940; Schechter & Lucero, 1968; Pentreath, 1971; Uchida & Irimura, 1974). The stomach consists of a single sac radially dividing into 10 pouches: five radial (ambulacral) pouches and five interradial (interambulacral) pouches (Pentreath, 1969). We also read a contradicting statement that brittle stars’ stomach totally has 15 swellings in a common context (Uchida & Irimura, 1974). Interradial pouches are well-folded structure deeply lying between the bases of arms, whereas radial pouches are limited in narrow spaces over arms (Pentreath, 1969, 1971; Uchida & Irimura, 1974). These pouches never extend into arms except the species *Ophiocanops fugiens* Koehler, 1922 (Fell, 1963).

Over and above anatomical studies, a dynamic perspective of digestion has been discussed based on live observation. In the green brittle star *Ophiarachna incrassata* (Lamarck, 1816), Wakita et al. (2018) reported a rhythmic movement termed “*pumping*” (Video S1; c.f. Fig. 1), which is frequently observed at the disk after feeding. The series of expansion and shrinkage can be recognized as a sort of peristaltic movement of the stomach. Its unique coordinated patterns are explainable by assuming internal fluid flows, the way of which depends on the morphology of the disk. In particular, five-armed individuals make unsynchronized movements between five body parts, whereas those are well synchronized in a six-armed case—peculiar individual difference in brittle stars. Thus, in this phenomenon, the morphological structure of the stomach may be of great importance, although it is unclear how pumping transforms it.

**Fig. 1.**
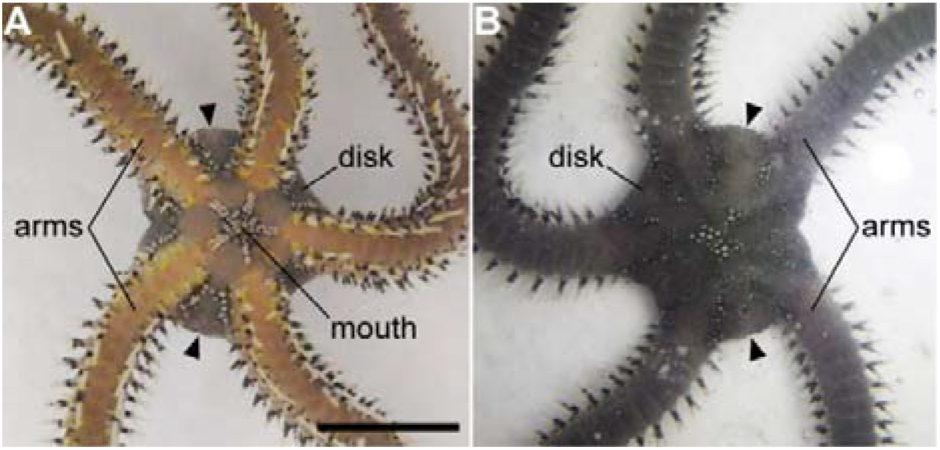
Frozen individual at the moment of the dynamic movement “pumping” in the green brittle star *Ophiarachna incrassata*. (A) Oral view. (B) Aboral view. Photos were taken just after pouring −80°C ethanol while we observed pumping at the disk. Arrowheads denote well-expanding portions. Scale bars represent 10 mm. Scanned and segmented images of this specimen are shown in Fig. 3. Video of another individual’s pumping is shown in Video S1.

In previous studies, the morphological structure of the digestive system in brittle stars has been visualized only by two-dimensional sketches or tissue sections (Smith, 1940; Pentreath, 1969, 1971; Schechter & Lucero, 1968; Uchida & Irimura, 1974; Frolova & Dolmatov, 2006, 2010). Moreover, traditional anatomy has employed a wide dissection of the body wall, which considerably distorts the morphology in focus. This issue arises from the fragileness of digestive organs as well as its deep attachment to the body wall by collagen strands (Schechter & Lucero, 1968; Uchida & Irimura, 1974). The aim of our study is to visualize non-destructive and three-dimensional (3D) morphological structure of the digestive system in brittle stars, in comparative terms of a relaxed condition and a condition during pumping. For this purpose, we employ X-ray micro-computed tomography (micro-CT) while introducing an instant freezing method using cryogenic ethanol for making a *snapshot* of the dynamically moving body (Fig. 1). We also probe into the internal morphology of the six-armed specimen studied by Wakita et al. (2018), so as to internally validate the prior assumption that this specific individual has six symmetrical units—previously it was made up merely in external terms. The primary conclusion in our study is that pumping could transform the entire stomach to help digestion in large brittle stars, with its coordinated patterns apparently reflecting five- or six-fold symmetrical arrangement in internal morphology.

## Methods

### Animals

Individuals of the green brittle star *Ophiarachna incrassata* were obtained commercially (Aqua Shop Saien, Sapporo, Japan) and reared in aquariums (600 × 600 × 600 mm) filled with artificial seawater at 25–28°C with the salinity of 32–35%o (TetraMarin Salt Pro, Tetra Japan Co, Tokyo, Japan). They were fed with dried krill (Tetra Krill-E, Tetra Japan Co, Tokyo, Japan).

### X-ray micro-computed tomography

With the aim to investigate relaxed and pumping bodies as well as five- and six-armed bodies, we chose three individuals: (1) a five-armed individual a day after feeding, with the disk diameter of 15 mm in a living anesthetized condition; (2) a five-armed individual a week after feeding, with the disk diameter of 18 mm; (3) a six-armed individual with the diameter of 25 mm, which had hardly shown feeding behavior for a few months. The six-armed one corresponds to that studied by Wakita et al. (2018). For the relaxed case, the animals (1) and (3) each were anaesthetized in 3% MgCl2 solution for an hour at room temperature and then fixed with Bouin solution—(1) for 10 days and (3) for five months—at 3°C with their arms cut near the bases. For looking into the pumping body, the animal (2) was put in a styrofoam box (127 × 157 × 100 mm) with 100–200 ml of artificial seawater and then fed with a dried krill. While we observed the rhythmic movement (pumping; c.f. Video S1), −80°C ethanol was poured onto the disk so that the animal (2) kept a momentary shape with expansion and shrinkage (Fig. 1). After slowly shaking the box for 10 min, the sample was put in Bouin solution for a day at 3°C with their arms cut.

After fixation, all the samples (1)–(3) were dehydrated with ethanol series (70%, 80%, and 90%) for two days each, and stained with 1% iodine diluted in ethanol for three days at 3°C to enhance the contrast of tissues in later X-ray exposure (Metscher, 2009). They were rinsed with 100% ethanol for a day at room temperature and then moved into *t*-butyl alcohol liquidized with a water bath above 40°C. After immersion in t-butyl alcohol for a day twice at 26°C, samples were superficially dried on tissues for several seconds and then put at −20°C for 10 min so that instantly frozen t-butyl alcohol would keep the original morphology as possible. They were freeze-dried by using a vacuum evaporator (PX-52, Yamato Ltd., Japan) with a cold alcohol trap (H2SO5, AS ONE, Japan). All chemicals were purchased from Kanto Chemical Co. (Tokyo, Japan).

Samples were scanned on an X-ray micro-CT system (inspeXio, SMX-100CT, Shimadzu Corporation, Kyoto, Japan), where X-ray source was operated at 75 kV and 40 μA. Scanned images were reconstructed and rendered by using VGStudio MAX ver. 2.2.6 (Volume Graphics, Heidelberg, Germany) with the voxel size of 10–50 μm. For segmentation of each sample using Amira ver. 2019.1 (Thermo Scientific, Waltham, USA), we traced the inner surface—boundary with contrasting X-ray absorptivity—of the digestive cavity beginning from the mouth. Note that segmentation was not available in regions where the inner wall stuck to each other so that the cavity was too flat to be identified. We also segmented gut contents, which were recognizable as highly absorptive areas inside the cavity. 3D animations were created with VGStudio MAX for slice images and Amira for segmented images, which are given in Videos S2–4.

## Results

The morphology of the digestive system, particularly the stomach, was well visualized by segmenting the inner wall of the digestive cavity (Figs 2–4). In the five-armed specimen fixed after anesthesia (Video S2), skeletal structure comprised five symmetrical sectors in appearance (Fig. 2A). The cavity’s surface was smoothly defined from the mouth to the stomach (Fig. 2B), so we required less subjectivity in the segmentation (Fig. 2C–E). The stomach comprised five larger interradial pouches and five smaller radial pouches (Fig. 2C). As noted in Methods, no region in a totally flat cavity was segmented, hence the gaps forming a cobweb-like structure could be interpreted as the missing flattened parts of the stomach, not representing there were many holes in morphology (Fig. 2C). The stomach was plain in the aboral side (Fig. 2C,D) but well wrinkled in the oral (Fig. 2E). Viewed orally, a ridge could be traced along each midline of two sorts of pouches, which descended into several branches (Fig. 2E). Interradial pouches narrowed at the bases with their breadth increasing distally, with the end being round so that we could trace smoothly between the oral and aboral surfaces (Fig. 2C–E). Radial pouches were more flat, shaped along arm skeletal plates, and made distal ends with rough and sharp edges (Fig. 2D,E). The distal parts of interradial pouches extended until near the oral wall, whereas their bases and radial pouches were restricted aborally (Fig. 2D). The oral room at the center contained a jaw apparatus and its peripheral organs, which would include the circumoral nerve ring and the water vascular system (Fig. 2A,B). In this specimen, two large pieces of food were respectively observed at the aboral base of an interradial pouch—might be partially shared by one adjacent radial pouch—and the oral end of another interradial (Fig. 2C–E).

**Fig. 2.**
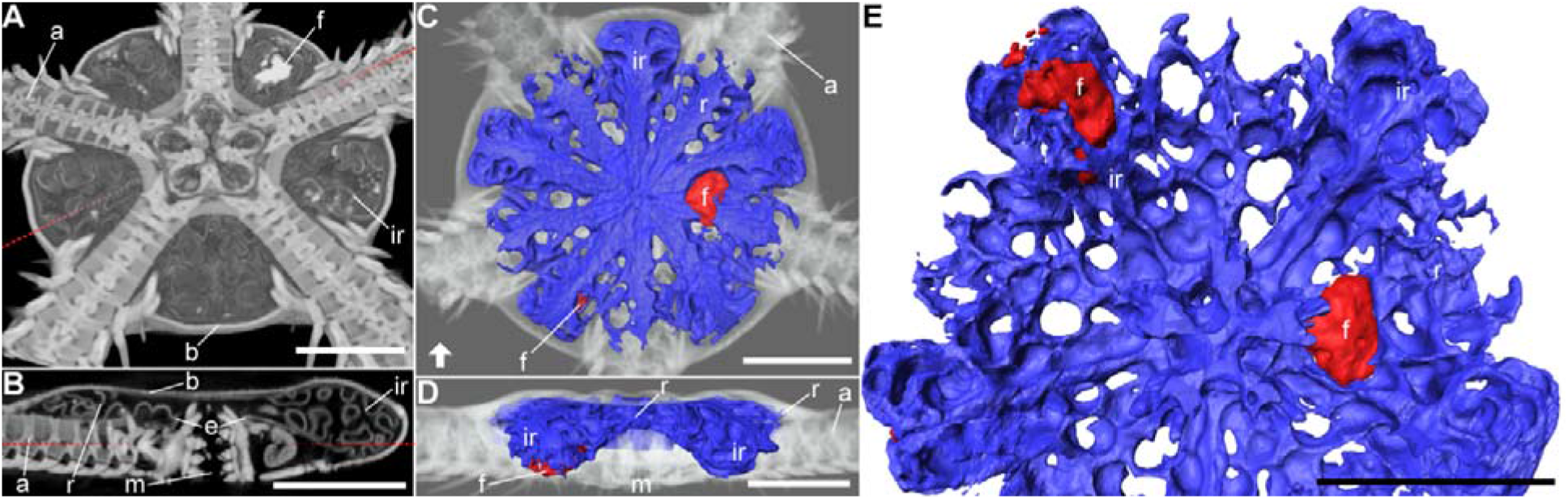
Three-dimensional visualization of the digestive system in a five-armed relaxed individual of the green brittle star *Ophiarachna incrassata.* Body structure was scanned with X-ray micro-computed tomography (micro-CT) and reconstructed in three dimensions, which is displayed in grayscale. The inner surface of the digestive cavity beginning from the mouth is colored blue. Contents in the stomach are colored red. Regions where the inward cavity was totally flat were not segmented (not colored blue) with a technical limitation, which reflects the apparent holes of the stomach and the apparent exposure of gut contents. (A) Reconstructed images viewed from the oral side, sectioned at a plane shown in (B) by the dotted line. (B) Oral-aboral section on a plane indicated in (A) by the dotted line; the bottom is oral; slab thickness is 0.38 μm. (C) Aboral view of the segmented model. (D) Lateral view of the segmented model from the side indicated in (C) by the arrow; the bottom is oral; the grayscale images are truncated in the front for clarity. (E) Enlarged oral view of the segmented model. Abbreviations: a, arm skeleton; b, body wall; e, esophagus; f, food (gut content); m, mouth; ir, interradial pouch; r, radial pouch. Scale bars represent 5 mm. Three-dimensional animation is shown in Video S2.

In the specimen frozen during pumping (Video S3), we found no noticeable damage due to the instant freezing method in external and internal morphology (Figs 1 and 3A). Besides, there seemed to be no large difference in the texture of the digestive wall in scanned images, compared to those observed in the relaxed one. We recognized a well-defined separation between the mouth and the stomach and a tight closure of the mouth (Fig. 3B; see the esophagus “e” and the mouth “m”), so we did not clearly identify the continuous space from the mouth opening. However, we had no ambiguity in segmenting the internal surface which was recognizable as the stomach’s one (Fig. 3C–E) when comparing to the relaxed case (Fig. 2C–E). The stomach wall during pumping was smoothly fitted to the distorted body wall with less obvious folding (Fig. 3C). The globular shaping of the stomach was also upheld from the observation that we saw no network-like segmentation denoting flattened patches (Fig. 3C), contrasting to the relaxed one (Fig. 2C). Expansions were remarkable in interradial pouches, which directed toward the aboral and lateral sides (Fig. 3D). Meanwhile, radial pouches at their neighbors also became more or less open (Fig. 3D). The oral surface likewise showed an entire extension, where we barely found sharp structure such as branched ridges (Fig. 3E). The cavity between the body and stomach walls (perivisceral coelom) appeared to be narrowly limited (Fig. 3B,D), not largely differing from the relaxed case (Fig. 2B,D). A food lied almost at the center of the stomach in this specimen (Fig. 3C,E).

**Fig. 3.**
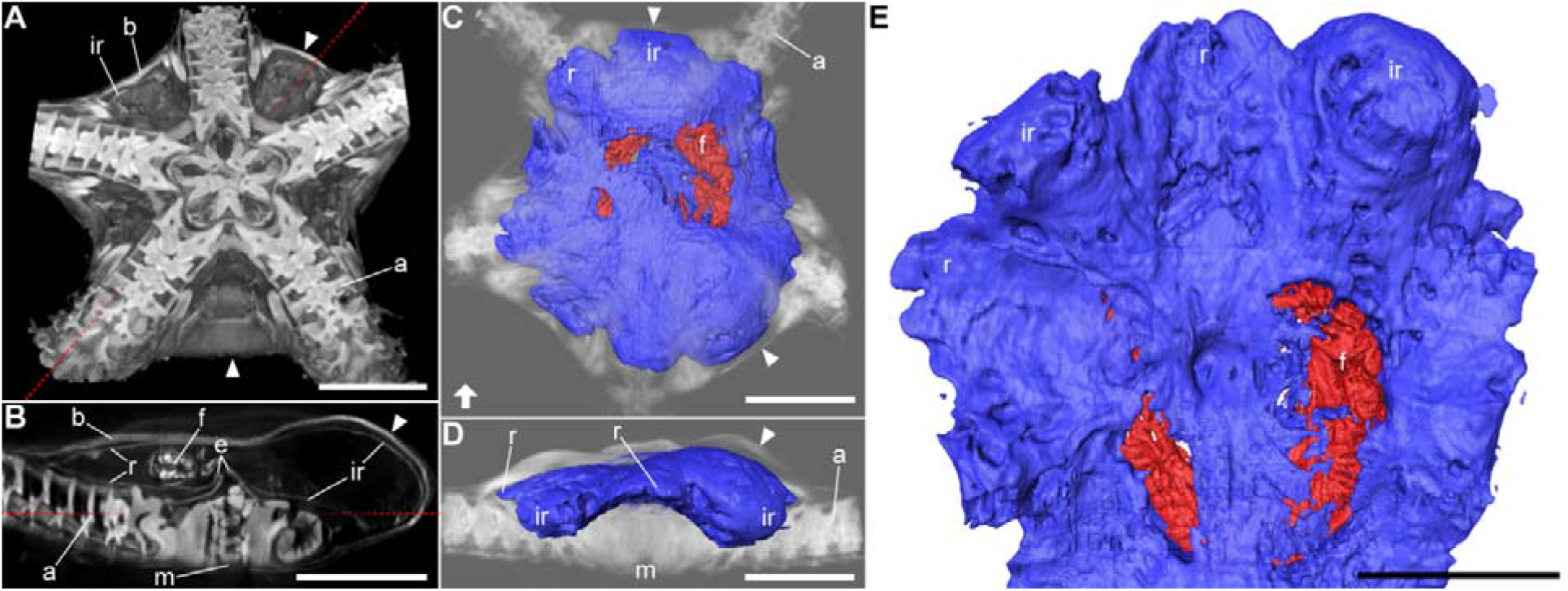
Three-dimensional visualization of the digestive system in a five-armed individual during the dynamic movement “pumping” in the green brittle star *Ophiarachna incrassata.* Body structure of an instantly frozen individual shown in Fig. 1 was scanned with X-ray micro-computed tomography (micro-CT) and reconstructed in three dimensions, which is displayed in grayscale. The inner wall of the digestive cavity is colored blue, which was identified in comparison with the relaxed specimen (Fig. 2). Contents in the stomach are colored red. Regions where the inward cavity was totally flat were not segmented (not colored blue) with a technical limitation, which reflects the apparent exposure of gut contents. Arrowheads denote well-expanding portions. (A) Reconstructed images viewed from the oral side, sectioned at a plane shown in (B) by the dotted line. (B) Oral-aboral section on a plane indicated in (A) by the dotted line; the bottom is oral; slab thickness is 0.36 μm. (C) Aboral view of the segmented model. (D) Lateral view of the segmented model from the side indicated in (C) by the arrow; the bottom is oral; the grayscale images are truncated in the front for clarity. (E) Enlarged oral view of the segmented model. Abbreviations: a, arm skeleton; b, body wall; e, esophagus; f, food (gut content); m, mouth; ir, interradial pouch; r, radial pouch. Scale bars represent 5 mm. Three-dimensional animation is shown in Video S3.

In the specimen with six arms (Video S4), a jaw apparatus and arm skeletal plates apparently arranged in six-fold radial symmetry (Fig. 4A). As in the five-armed relaxed case, the digestive cavity was smoothly traceable from the mouth to the stomach (Fig. 4B). Though the segmentation, we realized that the inner openings of interradial pouches were almost as flat as those of radial ones (Fig. 4B). This reduction was probably because this individual had not fed for a few months before fixation. Although the boundaries between radial and interradial pouches were less conspicuous (Fig. 4C) than the five-armed ones (Fig. 2C), the two types could be distinguished when we saw the model from several angles (Fig. 4B,C). In particular, interradial pouches hung down more orally (Fig. 4B) and showed ridged midlines on the oral surface (Fig. 4C). Here we could count 12 pouches—six interradial and six radial pouches (Fig. 4C).

**Fig. 4.**
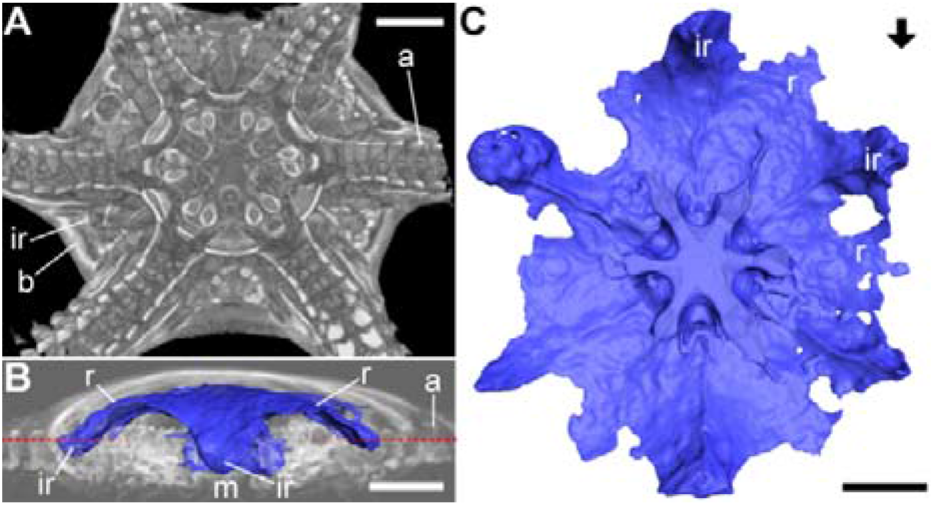
Three-dimensional visualization of the digestive system in a six-armed relaxed individual of the green brittle star *Ophiarachna incrassata.* Body structure was scanned with X-ray micro-computed tomography (micro-CT) and reconstructed in three dimensions, which is displayed in grayscale. The inner wall of the digestive cavity beginning from the mouth is colored blue. Regions where the inward cavity was totally flat were not segmented (not colored blue) with a technical limitation. (A) Reconstructed images viewed from the oral side, sectioned at a plane indicated in (B) by the dotted line. (B) Lateral view of the segmented model from the side indicated in (C) by the arrow; the bottom is oral; the grayscale images are truncated in the front for clarity. (C) Oral view of the segmented model. Abbreviations: a, arm skeleton; b, body wall; ir, interradial pouch; r, radial pouch. Scale bars represent 5 mm. Three-dimensional animation is shown in Video S4.

## Discussion

Our study has three main achievements. The first is 3D visualization of the uninjured stomach in brittle stars (Fig. 2, Video S2). The schematics were directly reconstructed from micro-CT scanned images, including less imagination than previous sketches. The second is introduction of a fresh methodology in micro-CT scanning, where we investigated a moving body’s momentary shape made by instant freezing with cryogenic ethanol (Figs 1 and 3, Video S3). The resultant snapshot provides a further understanding of the previously reported phenomenon, *pumping* (Wakita et al. 2018), as a possible digestive activity. The last is internal inspection of the specific six-armed individual studied by Wakita et al. (2018) (Fig. 4, Video S4). A supernumerary seemed to be simply a member of six equivalents, which helps a prebuilt assumption on pumping coordination.

In a five-armed representative, we counted 10 major swellings in the stomach (Fig. 2C) as Pentreath (1969) reported, not supporting another literature writing the total number 15 (Uchida & Irimura, 1974). Since the latter also mentions radial and interradial pouches, it might be miswriting or another interpretation given the complex folding. The wrinkles running on the oral surface would be a reflection in part of skeletal morphology and other organs’ anchoring, while probably being capable of a large amount of food by its extension. In fact, the specimen shown in Fig. 3 accommodates a relatively large food ranging across several pouches. Although the segmented stomach structure in Fig. 2 looks like a web network, we can interpret that these holes as untraceable flat parts of the cavity so that pouches connect with each other throughout the central space.

Comparison between the relaxed and pumping specimens in scanned images (Figs 2 and 3) indicates that pumping dynamically transforms the stomach, where it is natural to suppose internal fluid flows to some extent. The outer space of the stomach (perivisceral coelom) would make insignificant flows for pumping, considering the constant narrowness even in the pumping body (Fig. 3B,D). The mouth-stomach separation emerging in the pumping specimen (Fig. 3B) can be defined as the constriction of esophagus, referring to Schechter & Lucero (1968). This observation and the tight closure of the jaw apparatus—surrounding buccal cavity—(Fig. 3B) both would reinforce Wakita et al.’s (2018) assumption that the total fluid volume is constant during pumping; there is no outward leakage. Although their study built a water-connecting network with five nodes for the five-armed case, there could be 10 rooms given the number of pouches (Fig. 2C). However, the slits between the pouches were actually not deep as depicted in the pumping specimen (Fig. 3C). With this texture, the five-fold arm skeleton would rather work as partitions (Fig. 3A,B), making it reasonable to represent five rooms influencing the major behavior of internal flows. This explanation could also apply to the six-armed case. The rigid skeletal partitions symmetrically made by six arms (Fig. 4A) would be more dominant in fluid flow, compared to the flexibly transformable stomach with 12 pouches (Fig. 4C). Our scan also supports Wakita et al.’s (2018) explanation with six symmetrical nodes for this six-armed specimen. We thus retain the explanatory power of the pumping network with five or six symmetrical nodes—not 10 or 12 nodes in two different sizes.

Visibility in the original position of gut contents gives two suggestions for the phenomenon pumping. The first is about its initiation. After a brittle star eats something, food fragments would be seated at some pouches (Fig. 2C–E); even if a prey stays at the center, its body shape would never weight equally among all the pouches (Fig. 3C). The contents thus make the stomach morphology more asymmetric, which might trigger the initiation of a pumping series as Wakita et al. (2018) let one interradial volume unequal at the beginning of simulation. The second involves the purpose of pumping. In many animals including humans, food transfers from the mouth to the anus in one direction; the linear structure guarantees that nutrients are absorbed point by point. On the other hand, the digestive cavity of brittle stars has neither unidirectional tracts nor the anus, so a piece might easily stick to a dead end—just as shown in Fig. 2. Transformation of the stomach by pumping would give more opportunities to displace the piece with it spreading a nutritious flow. This strategy would be effective in large-sized species, where gut contents travel longer distances piece by piece. Therefore, although we used the single species *Ophiarachna incrassata,* other large brittle stars are supposed to exhibit pumping in a similar manner.

CT scanning technique for 3D visualization has been employed by several studies on brittle stars. Landschoff & Griffiths (2015) revealed how a brooding brittle star accommodates several juveniles inside its body, comparing two species; another was similarly examined later (MacKinnon et al. 2017). In a taxonomic context, Okanishi et al. (2017) described skeletal structure of a euryalid brittle star without dissolving its thick skin. Clark et al. (2018) paid attention to the joint connection of vertebrae to understand the mobility of arms in two species. Our study would carry a novelty in (1) focusing on the digestive system, (2) comparing two behavioral conditions (relaxed v.s. pumping), and (3) comparing two morphologically different individuals within a species (five-armed v.s. six-armed) in CT scanned brittle stars. Notably, the snapshotting method for (2), where cryogenic ethanol is poured onto a living animal, is widely applicable for the scanning purpose of body structure during dynamic movements in echinoderms. These approaches give prominence to a dynamic digestive process and its relation with body structure, which would be a hot clue to ethology, ecology, and evolution in echinoderms.

## Acknowledgments

This work was partly supported by JSPS KAKENHI (Grant Number 16KT0099), JST CREST (Grant Number JPMJCR14D5), and Hokkaido University Frontier Foundation (Nitobe School Financial Assistance), Japan.

## Author contributions

D.W. and H.A. designed the study, D.W. conducted experiments, H.A. performed micro-computed tomography, K.N. conducted segmentation, D.W. drafted the manuscript and prepared figures and videos, K.N. and H.A. revised the manuscript, and all the authors approved the article.

## Supporting Information

**Video S1**. Rhythmic movement “pumping” in a five-armed individual of the green brittle star *Ophiarachna incrassata.*

**Video S2**. 3D animation of the images shown in Fig. 2 (five-armed relaxed individual).

**Video S3**. 3D animation of the images shown in Fig. 3 (five-armed pumping individual).

**Video S4**. 3D animation of the images shown in Fig. 4 (six-armed relaxed individual).

